# Using protein language models for pangenome construction

**DOI:** 10.64898/2026.06.04.730042

**Authors:** Niels Jakob Larsen, Pep Charusanti, Henry Webel, Louis Ohl, Kai Blin, Jes Frellsen

## Abstract

Current pangenome construction methods rely largely on nucleotide or protein sequence alignment, limiting their ability to detect remote orthologs and semantic relations. We introduce a novel method that leverages protein language model embeddings to capture functional and semantic relationships beyond sequence similarity. Our approach employs approximate nearest-neighbor search coupled with a clustering step utilizing HDBSCAN, DBSCAN, or weighted single-linkage clustering with multiple similarity thresholds. The method utilizes GPU acceleration, dynamic batching, and ONNX optimization to scale approximately linearly with the number of proteins, enabling the analysis of datasets containing millions of proteins. We evaluated our approach on a randomly sampled subset of OrthoDB and the CAFA5 dataset, benchmarking it against SCARAP. SCARAP is a recently published tool with similar performance to a variety of other common tools for computing pangenomics. Our benchmarking demonstrates that our method produces more specific clusters than SCARAP across both datasets. SCARAP excelled in term consistency within clusters on the OrthoDB dataset, where labels are inferred with sequence alignment (using MMseqs2). Both methods face a significant degradation in term consistency when transitioning to the experimentally validated CAFA5 dataset, ultimately resulting in similar term consistency scores for both approaches. Crucially, our approach yields superior cluster quality on both datasets and significantly outperforms SCARAP across all metrics of functional consistency and coherence on the experimental CAFA5 dataset. Finally, we demonstrate the method’s scalability and utility by characterizing the pangenome of 1,034 Streptomyces genomes. The pipeline is available for use at our GitHub: https://github.com/jakob949/pan_genome

## 1 Introduction

Pangenomics is the study of the complete set of genes found within a defined group of genomes. This is the idea of moving beyond a single reference genome to representing the full spectrum of genetic diversity within a group of genomes. This concept has enabled pangenomics methods to give a better understanding of genetic diversity, evolutionary dynamics, the functional implications of genomic variation, and niche adaptation [1]. Classically, the pangenome is structured into three subgroups: The core genome consists of genes found in almost every genome, which represents the fundamental genetic parts. The shell genome includes genes present in most but not all genomes, reflecting common functions and adaptations. Finally, the cloud genome is composed of genes unique or almost unique to a single genome, therefore showing strain-specific traits. Analyzing the distribution of genes across these groups (as illustrated in Table 1) allows one to uncover the genetic basis for specific traits, possible diseases, and environmental adaptations [1, 2].

**Table 1.** Genome proportion thresholds used to classify protein clusters into core, shell, and cloud pangenomic classes.

| Gene Group Class | Frequency in Genomes |
| --- | --- |
| Core | > 95% |
| Shell | [15%, 95%] |
| Cloud | < 15% |

**Table 2.** Performance comparison on the OrthoDB dataset. The best performance for each metric is highlighted in bold. Default settings were used for SCARAP. Parameters for the weighted multi-threshold single-linkage clustering are model-specific: for ProtT5, mean = 0.856 and SD = 0.1259; for ESM2 3B, mean = 0.9935 and SD = 0.0076. For density-based clustering ((H)DBSCAN), parameters were set to ϵ = 0.045 and minimum cluster size = 2 for both the ProtT5 and ESM2 3B models. FC denotes Functional Consistency.

| Metric | ProtT5 Embeddings |  |  | ESM2 3B Embeddings |  |  | SCARAP |
| --- | --- | --- | --- | --- | --- | --- | --- |
|  | Multi Threshold | DBSCAN | HDBSCAN | Multi Threshold | DBSCAN | HDBSCAN |  |
| Mean Hierarchical Purity | 17.6714 | **18.613** | 15.6819 | 15.9472 | 16.4352 | 14.7595 | 11.2521 |
| Weighted Mean Hierarchical Purity | 11.9043 | **14.9876** | 13.4053 | 6.0949 | 10.4948 | 10.0196 | 6.06 |
| Mean GO Coherence | 5.5577 | **6.3638** | 5.74 | 5.1952 | 5.1906 | 5.0106 | 3.5855 |
| Weighted Mean GO Coherence | 5.5786 | **7.2994** | 6.532 | 3.3387 | 5.8094 | 5.6642 | 4.2484 |
| Mean FC (%) | 97.2658 | **96.894** | 89.6619 | 99.8171 | 93.0385 | 87.9247 | 84.0109 |
| Weighted Mean FC (%) | 81.6556 | **88.7844** | 81.082 | 45.9251 | 73.6911 | 71.122 | 62.6711 |
| Term Consistency Score | 0.3153 | 0.279 | 0.3486 | 0.6327 | 0.5108 | 0.5573 | **0.8731** |

The variability of pangenomes intuitively stems from the different sets of genomes being compared. A primary way of characterizing this variability is to distinguishing between “open” or “closed” pangenomes. In an open pangenome, sequencing of additional genomes continues to reveal new genes, demonstrating an expanding gene pool. Conversely, a closed pangenome reflects a complete gene set, sequencing additional genomes yields few novel genes. Factors such as population size, evolutionary rate, and the variety of ecological niches occupied by a genotype ultimately drive this degree of openness [3]. The balance of gene acquisition, mainly through horizontal gene transfer, and gene loss is what gives rise to pangenomes. The evolutionary processes at work produce the genetic diversity required for an organism to survive different environments and adapt to various ecological pressures [4]. This is a main focus of pangenome research.

Pangenome analysis is essential for evolutionary research, metagenomics, comparative genomics, and natural product discovery. Since the maturation of the concept of pangenomics two decades ago [5], several methods, software packages, and pipelines have been developed to compute the pangenome. Some of the best-known approaches include Roary [6], Panseq [7], PanX [8], anvi’o [9], and SCARAP [2]. These methods typically rely on sequence alignment, k-mer comparisons, or various clustering techniques [10] and recently graph based approaches are emerging [11]. All the tools have the same underlying concept of clustering ortholog sequences from the input genomes. A core challenge in clustering ortholog sequences, particularly for diverse species sets, such as open pangenomes lie in accurately identifying orthologous genes, especially those that are distantly related. Traditional sequence alignment methods often struggle in the “twilight zone” of amino acid sequence identity (less than 30%), giving rise to misclassification when doing pangenomics analysis of distant genes [12].

Advances in deep learning, specifically Protein Language Models (pLMs) like ESM variants [13] and ProtTrans [14], offer a powerful alternative. These models, typically based on the transformer architecture, are trained on massive unlabeled sequence datasets via self-supervised masked language/token modeling (MLM training). The core transformer mechanism is attention, which captures complex, long-range dependencies within sequences by assigning weights to each input token (e.g., amino acid) and computing a weighted sum of value vectors, allowing the model to focus on relevant information [15]. As visualized in Figure 1, amino acids are tokenized, assigned positional encodings, and passed through *N* stacks of attention layers and non-linear fully connected neurons, producing an embedding for each input token (amino acid). A fixed-dimensional protein embedding is often obtained by averaging these amino acid embeddings. This process generates dense vector representations called embeddings that capture intricate patterns related to protein structure, function, and evolution, going beyond sequence similarity [16, 17]. This information-rich encoding captures complex sequence dependencies and structural cues often overlooked by conventional methods, allowing for improved detection of remote orthologs. Because protein structure diverges more slowly than sequence, these embeddings can implicitly encode structural information, enabling pLM-based methods to excel at remote homology detection. Tools like pLMSearch [17] and TM-Vec [16] have demonstrated that embeddings can identify orthologs which have dissimilar sequences but with similar protein structure, increasing sensitivity compared to methods like BLASTp or MMseqs2 [17]. Those tools are approaching the accuracy of direct protein structure comparison without requiring 3D coordinates. However, a critical consideration is the computational cost of classical attention, where calculating the attention matrix involves operations over all token pairs, leading to O(n^2^) time complexity, where *n* refers to the sequence length.

**Fig. 1.**
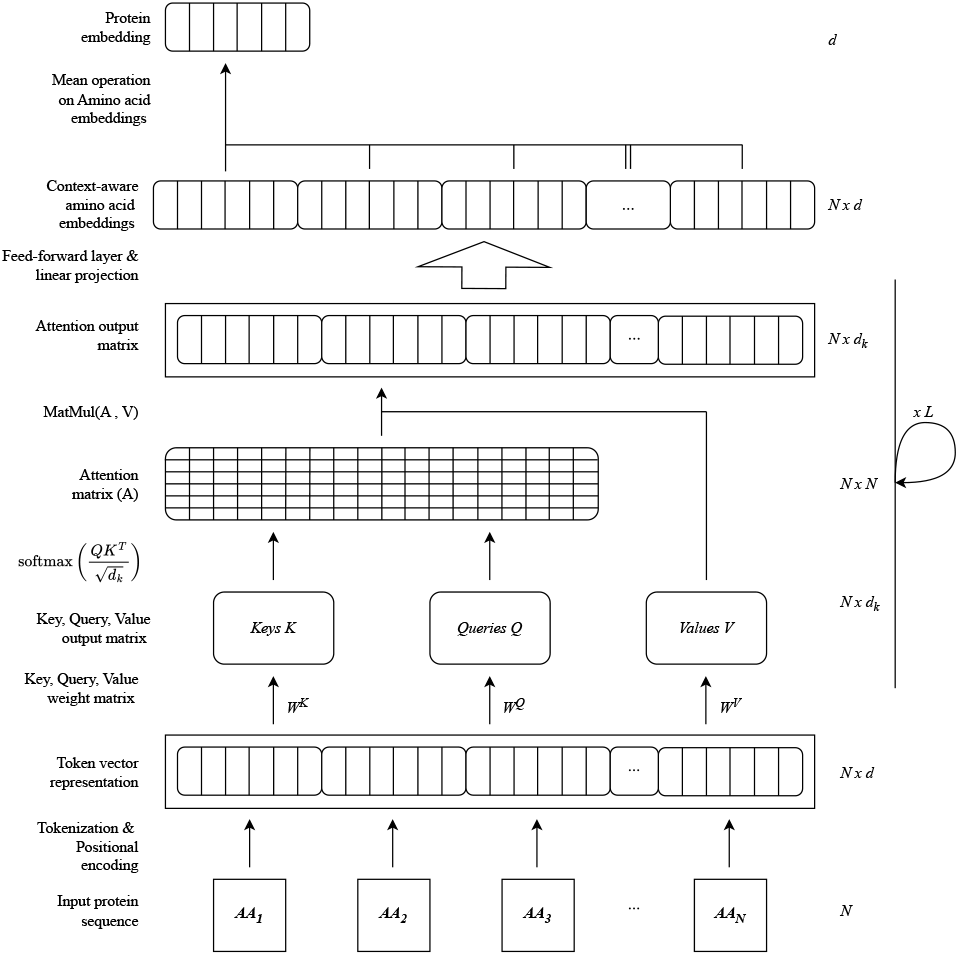
Schematic of a Transformer-based protein sequence encoder. The architecture processes an input sequence of *N* amino acids (AA) through tokenization and positional encoding to generate token vector representations of dimension *N × d*. The attention mechanism computes Queries (Q), Keys (K), and Values (V) via linear projections to generate an *NN* attention matrix *A* using scaled dot-product attention. The resulting attention output (*Ndk*) undergoes a feed-forward operation and linear projection to produce context-aware embeddings of size *Nd*. This process is repeated for *L* layers. A final mean pooling operation aggregates the sequence into a fixed-length protein embedding of dimension *d*. To simplify the figure is residual connections and layer normalization not displayed.

**Fig. 2.**
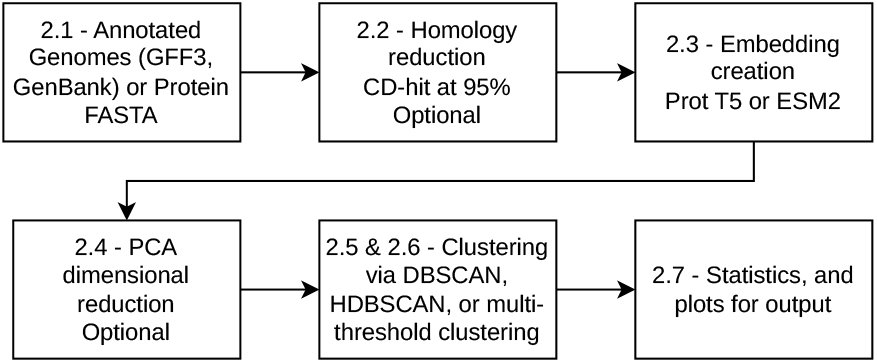
Flowchart describing the overall workflow of the project. The numbers in the figure refer to the subsection numbers.

The ability of pLMs to capture distant relationships is highly relevant for pangenome construction. Accurately identifying remote orthologs is essential for defining robust core genomes, understanding the evolution of accessory gene families, and tracing horizontal gene transfer events. Even though pLMs have shown success in homology search (pLMSearch [17], TM-Vec [16]), functional annotation, and their foundational role in structure prediction tools like AlphaFold [18] and ESMFold [19], no existing pangenome construction method utilizes these information rich protein embeddings to compute the core, shell, and cloud subgroups. As mentioned, current tools rely primarily on explicit sequence-level comparison. This represents a missed opportunity to utilize the nuanced understanding of protein relationships captured by pLMs.

In this work, we present a novel scalable pangenome method that uses embeddings from protein language model as the primary basis for clustering proteins, thereby harnessing the information rich embedding for improved sensitivity for pangenome construction. The pipeline is available at GitHub https://github.com/jakob949/pan_genome

## 2 Methods

The method combines established bioinformatics and deep learning methods with novel approaches for fast clustering of embeddings. Our implementation leverages GPU acceleration for all computationally intensive steps and achieves empirically near-linear scaling with respect to the number of total genes in the analysis. Our method excels in producing large-scale pangenomics analysis across distant species.

### 2.1. File parsing

The primary purpose of the file parsing step is to extract the complete set of translated protein sequences from the genomes intended for pangenome analysis. The input parser accepts multiple files in FASTA, GenBank, or GFF3 formats. The pipeline treats the sequences contained within each distinct file as belonging to the same genome, contig, or chromosome, assigning a shared genome ID based on the respective filenames.

The pipeline does not compute the pangenome from unannotated whole-genome nucleotide sequences. The input files must contain pre-computed protein annotations. For GenBank and GFF3 formats, the parser isolates features annotated as coding sequences (CDS) and extracts their corresponding amino acid translations. For FASTA formats, the pipeline expects the amino acid sequences of the proteins to be formatted as individual sequence records within a single file per genome. Detailed specifications for input file formats are available in the repository documentation.

### 2.2. CD-hit (optional)

Following file parsing, an optional step of CD-HIT [20] performs homology reduction exclusive within each genome to reduce computational complexity in subsequent steps . We employ a sequence identity threshold of 95% to retain sequence diversity while eliminating near-identical sequences. These near-identical sequences will be treated as a single sequence for the later steps. We select a sequence of the near-identical sequences as the representing sequence.

### 2.3. Sequence embedding

To capture meaningful representations of protein sequences, we employ pretrained transformer-based encoder models with an architecture similar to that illustrated in Fig 1. Specifically, our pipeline integrates and utilizes two primary models: the ESM2 3B model [13] and the encoder portion of ProtT5-XL-UniRef50 [14]. Both models are utilized in this study as they are well-suited for capturing homology signals from diverse protein sequences. Each protein sequence is fed into the chosen encoder. The resulting protein embedding vectors possess a fixed dimensionality, specifically 1024 dimensions for the ProtT5-XL encoder and 2560 dimensions for ESM2 3B. This uniform dimensionality is maintained irrespective of the input sequence length via mean pooling across the token sequence, illustrated in the last step of Fig 1. This consistency in dimensionality ensures that our subsequent similarity-based clustering step can operate on any embedded protein set.

We implement the embedding step in PyTorch using Huggingface’ Transformers library. By default, the tool uses “ESM2 3B”. The computational intensity of creating embeddings requires GPU acceleration to have a feasible complete time (however, it is possible to use CPU). To enhance scalability and performance, we incorporate dynamic batching techniques. Additional performance optimization strategies, including model quantization and ONNX integration, are described in detail in the subsequent section: Accelerated embedding creation.

### 2.4. PCA (optional)

There is an optional setting to perform PCA dimensionality reduction along the embedding dimension. This is a recommended step when the dataset is very large to reduce the computational cost of the subsequent clustering step.

### 2.5. Clustering, and Approximate Nearest Neighbor

Central to our pangenome construction is the clustering of protein embeddings derived from pLM. Clustering in an embedding space allows grouping proteins based on learned features that can capture functional, structural similarities, or other rich features, beyond primary amino acid sequence identity [16, 17].

Naively, distance-based clustering is computationally expensive due to the necessity of all-against-all comparisons. To drastically reduce the number of comparisons, we employ approximate nearest neighbor search. Our method incorporates three options for clustering embeddings: weighted multiple thresholds single-linkage clustering, DBSCAN [21], and HDBSCAN [22]. These clustering algorithms and the approximate nearest neighbor search are explained in the subsequent sections. After approximate nearest neighbor search and clustering, each embedding is assigned a cluster ID and is assigned a pangenomic class: core, shell, or cloud using the frequency thresholds in Table 1 as defaults. The calculation of the assignment of cluster ID and pangenomic class are explained in Section 2.5.8.

#### 2.5.1. Approximate Nearest Neighbor

To allow for clustering of potentially millions of protein embeddings efficiently, we employ Approximate Nearest Neighbor (ANN) search using the FAISS library [23]. ANN significantly speeds up the process of clustering by only taking similar embeddings into consideration for building a given cluster. This is much faster compared to an exact, exhaustive search, all-against-all comparison.

The ANN search can be formulated as follows: given a reference matrix X ∈ ℝ^*N* ×*D*^, composed of N stacked individual protein embeddings x_*n*_ ∈ ℝ^*D*^ where D represents the embedding dimensionality, and a query vector y ∈ ℝ^*D*^ . The objective is to retrieve the k nearest embeddings to y from X *without* executing an exhaustive O(N ) search.

We ensure metric consistency by strictly L_2_-normalizing all embeddings in the reference matrix X and query vectors y prior to any indexing or search operations.

To avoid performing an exhaustive O(N ) scan, we utilize an Inverted File (IVF) index structure. This structure partitions the globally normalized reference matrix X into m Voronoi cells (k-means regions) defined by a set of centroids Π = {*µ*_1_, …, *µ*_*m*_} ⊂ ℝ^*D*^ . Because the data resides on a unit hypersphere, the Euclidean distance used to define these regions effectively maps to cosine similarity. Each normalized embedding x_*n*_ is assigned to the cell of its nearest centroid. Consequently, the search space is reduced from the full dataset X to a specific subset of candidate embeddings. This process of retrieving the k nearest vectors happens in two steps:

1. **Coarse Quantization:** First, the closest centroid *µ*^∗^ to the query y is determined by calculating the minimum distance to the set of centroids Π:

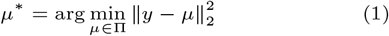
2. **Fine Search:** The exact similarity calculation is restricted to the subset of embeddings 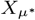 residing within the k-means region of *µ*^∗^. Because both the candidate vectors 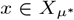 and the query vector y are already L_2_-normalized, the cosine distance is efficiently computed via the inner product:

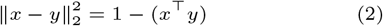

This approach approximates the nearest neighbors by assuming the true neighbors of y are located in the same (or adjacent) k-means regions, reducing the complexity significantly.

In practice, the ANN search is performed by the FAISS library, which queries the k + 1 nearest neighbors to account for the inclusion of the query vector itself. The returned cosine similarity scores are then transformed into cosine distances. For density-based clustering (HDBSCAN and DBSCAN), these neighbor distances are accumulated to construct a sparse k-NN distance graph, represented mathematically as an adjacency matrix 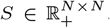 . Within this structure, only the identified neighbor relationships are explicitly instantiated as distance edges; non-neighbor entries remain structurally unallocated. For weighted multiple thresholds single-linkage clustering, the global matrix S is bypassed entirely, and the batch-level similarity arrays are fed directly into a Union-Find (disjoint-set) data structure on the fly, following the Algorithm 1 seen in appendix. The disjoint-set data structure efficiently tracks and merges non-overlapping subsets of proteins as new similarity edges are processed, allowing for the rapid assembly of connected components.

Using ANN involves a theoretical trade-off, accepting a small potential loss in accuracy for substantial gains in speed, which is essential for large-scale pangenomic analysis.

#### 2.5.2. Clustering methods used

The clustering phase proceeds via one of three user-selected algorithms: DBSCAN, HDBSCAN, or weighted multiple thresholds single-linkage clustering. For the density-based methods (DBSCAN and HDBSCAN), the sparse similarity Matrix 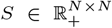 generated by the ANN search serves as the direct input. DBSCAN and HDBSCAN are implemented with the scikit-learn library. The weighted multiple thresholds single-linkage clustering bypasses the matrix S by processing the k nearest points directly into a Union-Find data structure on the fly, see Algorithm 1 seen in appendix.

#### 2.5.3. Density-Based Spatial Clustering of Applications with Noise (DBSCAN)

DBSCAN identifies clusters within the sparse similarity matrix S by evaluating local density [21]. The algorithm is parameterized by a minimum number of neighbors (minPts) set to 2 and a static distance boundary ϵ set to 0.05 for ESM2 3B and 0.045 for ProtT5-XL. A core point is defined as an embedding with at least minPts neighbors within the ϵ radius. Clusters expand by connecting mutually reachable core points and their associated border points. Embeddings that fail to meet this minimal reachability criterion are classified as noise, preventing the erroneous merging of unrelated sequences.

#### 2.5.4. Hierarchical DBSCAN (HDBSCAN)

HDBSCAN extends the DBSCAN paradigm by employing a mutual reachability distance metric to construct a hierarchical linkage tree, effectively evaluating clusters across all possible density thresholds [22]. While the algorithm primarily extracts a flat partition by maximizing cluster stability throughout this hierarchy, it also incorporates a minimum number of neighbors minPts parameter which is set to 2, and a cluster selection ϵ value is set to 0.05 for ESM2 3B and 0.045 for ProtT5-XL. In this implementation, ϵ serves as a lower-bound distance threshold, forcing to merge micro-clusters that fall below the specified distance to ensure more robust and biologically meaningful cluster definitions.

#### 2.5.5. Weighted multiple-threshold single-linkage clustering

The heterogeneity of orthologous group divergence is a critical factor in pangenome construction, as illustrated by the cluster consistency profiles in Figure 3A. When the protein sequences representing orthologous gene groups are embedded using ProtT5-XL and clustered via traditional single-linkage clustering at varying similarity thresholds (between 0.5 and 1.0), it effectively demonstrates the specific similarity thresholds at which different orthologous gene groups fragment. The ideal outcome for any pangenomic method is to assign all proteins from an orthologous gene group into a single cluster, meaning the similarity threshold should be positioned precisely before fragmentation occurs. However, the cluster fragmentation behavior observed in Figure 3B suggests that each orthologous group ideally requires a distinct similarity threshold. Therefore, it is evident that a single global similarity threshold fails to capture the evolutionary diversity across multiple ortholog gene groups.

**Fig. 3.**
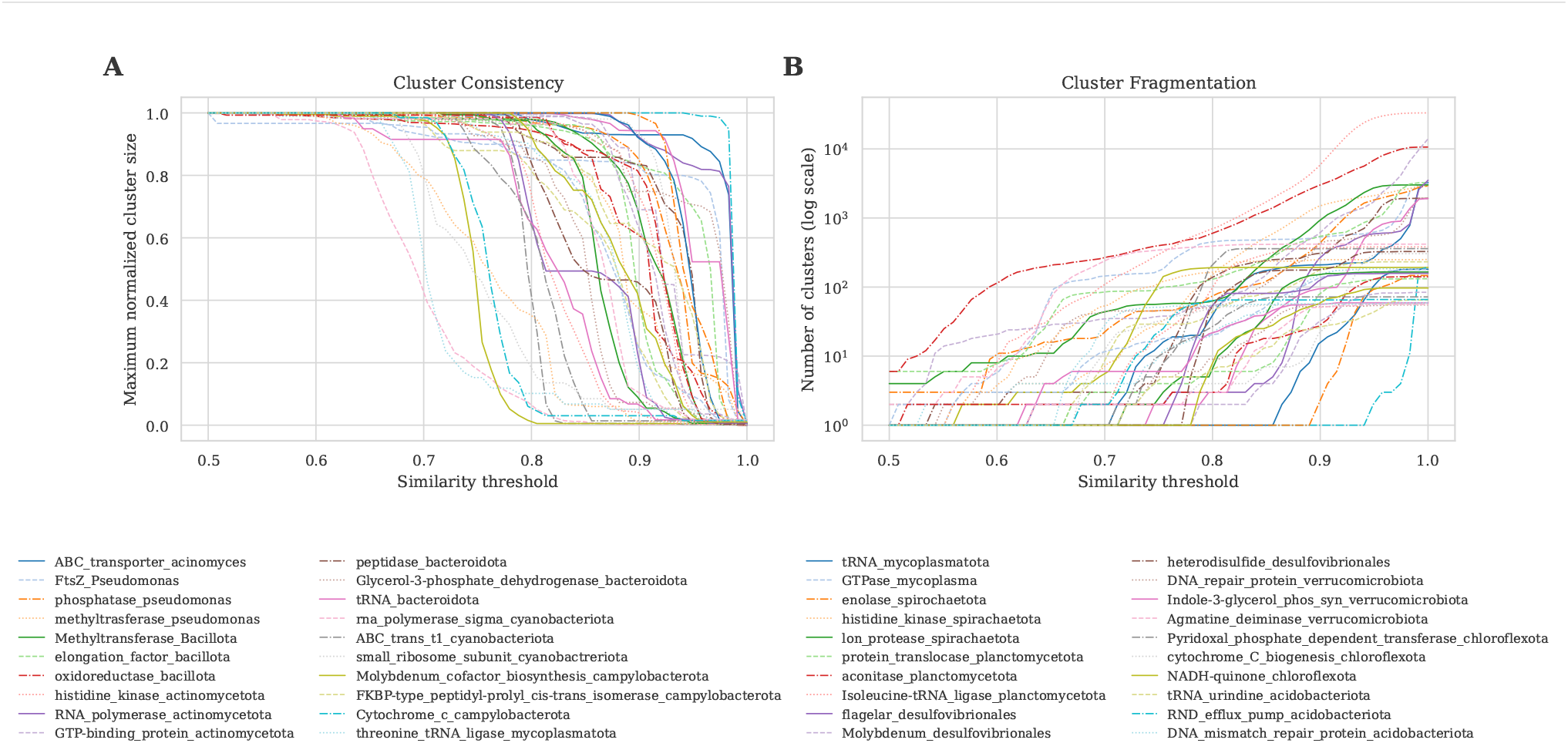
Impact of similarity thresholds on ortholog group cluster formation. Part A demonstrates how different similarity thresholds affect the size of the largest cluster for different ortholog groups. In Part A, a value of 1 maximum normalized cluster size means that all proteins from the ortholog group are contained within a single cluster. At those similarity thresholds that result in a y-value of 1 indicate that the clustering algorithm has successfully merged all ortholog protein embeddings into a single cluster. Conversely, a value close to 0 implies that the group has been partitioned into a greater number of smaller clusters or singletons. The important observation from Part A is that different ortholog gene groups require different similarity thresholds to be correctly classified as a single ortholog group. Part B illustrates how different ortholog gene clusters fragment at if a overall similarity threshold is used. It can be observed from Part B that cluster fragmentation occurs even at low similarity thresholds.

To address this limitation, weighted multiple-threshold single-linkage clustering executes multiple iterations of the single-linkage clustering algorithm, with each iteration utilizing a distinct distance threshold to generate different cluster formations. To derive a single consensus assignment from these disparate runs, we apply a probabilistic weighting scheme based on the concept of cluster “tipping points.” We define a tipping point as the distance threshold at which a complete orthologous gene group begins to fragment, causing its classification to drop from the “core” pangenome (e.g., ≥ 95% genome conservation) into the “shell” category. Because different orthologous groups tolerate different levels of divergence, these tipping points vary across gene families. By empirically evaluating 34 different orthologous gene groups, we observed that their tipping points form a normal distribution (illustrated in Figure 7 in the appendix). The same empirical idea was then expanded to include 380 otholog gene groups, and the mean and standard deviation for the distribution was found to be, for the ESM2-3B model, mean: 0.9935 and SD: 0.0076. For the ProtT5-XL, mean: 0.856 and the SD: 0.1259.

By using weighted multiple-threshold single-linkage clustering we do try to yield a higher biological meaning compared to traditional single-linkage clustering. The mechanism of cluster formation using this approach is detailed in Section 2.5.8 (Equations 4 and 5), and an implementation is provided in Algorithm 1 in the appendix. Finally, the application of multiple thresholds introduces a significant challenge regarding the meaningful evaluation of all resulting clusters, which is further addressed in Section 2.5.8.

#### 2.5.6. Avoiding intra-genome cluster expansion

For all three clustering approaches, to avoid the bias of cluster formation or expansion driven by protein embeddings originating from the same individual genome, edges connecting proteins encoded within the same genome (intra-genome protein pairs) are explicitly omitted from the sparse distance matrix S (or union-find structure for the weighted multiple-threshold single-linkage clustering). Thus, two intra-genome proteins can only be assigned to the same cluster on the condition that they share a transitive path through homologous genes from other individuals.

#### 2.5.7. Probabilistic Category Assignments for density clustering

After density clustering (DBSCAN or HDBSCAN), each protein classified as noise is assigned a unique cluster ID. This gives K clusters 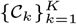, including a noise cluster.

To determine the pangenomic category of a cluster, we calculate the proportion of total genomes represented within it. Let N be the total number of genomes in the analysis, and let g(p) denote the genome from which protein p originates. For a given cluster C, the set of unique genomes contributing at least one protein to this cluster is given by {g(p) | p ∈ C}. The genome proportion f (C) is calculated as the size of this set divided by N :

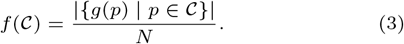

Pangenomic categories are then assigned based on the proportion thresholds defined in Table 1 or by the user.

#### 2.5.8. Probabilistic Category and Cluster Assignments for Multi-Threshold Single-Linkage Clustering

After applying multi-threshold single-linkage clustering, we retain the clustering results across the set of M similarity thresholds T = {τ_1_, τ_2_, …, τ_*M*_ }. To evaluate these results, we employ a weighted probability scheme to determine the consensus assignment for both pangenomic categories (Eq. 4) and cluster memberships (Eq. 5).

There are multiple ways to determine the weights, w_*j*_, corresponding to each threshold τ_*j*_ . While a baseline approach uses an unweighted majority vote (w_*j*_ = 1 for all j), a more rigorous method applies the normally distributed tipping points established earlier (Appendix Figure 7). Deriving w_*j*_ from this distribution directly aligns the clustering model with the empirical biological transition probabilities observed in ortholog groups.

To calculate the specific pangenomic category for a protein p, we compute a weighted probability across the threshold set. Let C_*p*_(τ_*j*_ ) denote the category assigned to protein p at threshold τ_*j*_, calculated using the genome proportion logic from Equation 3 and Table 1. The probability that protein p belongs to a specific consensus category C ∈ {Core, Shell, Cloud} is computed as:

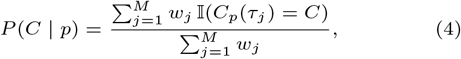

where 𝕀 is the indicator function yielding 1 if the condition is true and 0 otherwise. The final category for protein p, denoted 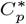, is the one that maximizes this probability: 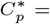 arg max_*C*_ P (C | p). The maximized value serves as a confidence score for the assignment, reflecting its consistency across the weighted thresholds.

A similar approach is used to determine the final cluster ID for each protein. Let k_*p*_(τ_*j*_ ) be the temporary cluster ID assigned to protein p at threshold τ_*j*_ . The weighted probability that protein p belongs to a specific cluster k across all the thresholds is computed as:

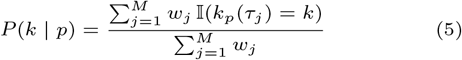

Protein p is then assigned the final cluster ID, K^∗^, that maximizes this aggregated probability (k^∗^ = arg max_*k*≤*K*_ P (k | p)). This probability indicates the confidence in the cluster assignment, reflecting how consistently the protein is grouped into cluster k^∗^ across the threshold set T .

The assignment of cluster ids is deterministic for multi-threshold single-linkage clustering. Because every protein p starts in its own singleton cluster (effectively its own index), the “Cluster ID” at any given threshold τ_*j*_ is simply the index of the root element in the Union-Find structure.

### 2.6. Compute acceleration and space optimizations

To facilitate the analysis of large-scale biological datasets, we optimized the most computationally intensive components of the pipeline: embedding generation and clustering, to ensure efficient scalability.

#### 2.6.1. Accelerated embedding creation

The generation of pLM embeddings represents the most computationally intensive component of the pipeline. While upstream homology reduction (see Section 2.2) minimizes the input volume, accelerating the embedding process itself is crucial for scalability. To facilitate efficient processing on hardware ranging from high-performance clusters to consumer-grade GPUs, we optimized the transformer inference for both memory usage and speed. The primary bottleneck—forward-pass computation—was addressed by implementing the ONNX runtime and INT8 quantization.

To evaluate the fidelity of these optimizations, we benchmarked them on a dataset of 9,000 genes. Embeddings generated by the optimized models (ONNX runtime and INT8 quantized) were compared against the baseline PyTorch implementation via pairwise cosine similarity. The results indicate that both optimizations yield representations virtually identical to the baseline, with mean cosine similarities (and standard deviations) of 1.0000 (0.0000) for the ONNX model and 0.9999 (0.0005) for the INT8 quantized model.

Performance benchmarks reveal that INT8 quantization reduced inference latency by 35%, while the ONNX runtime decreased total computation time by 70%. However, it is important to note that the ONNX runtime incurs an initialization overhead, which must be weighed against speed gains in scenarios involving small datasets or few repeated inferences. ONNX runtime is not compatible with ESM2 models.

#### 2.6.2. Dynamic batching

Dynamic Batching determines the batch size based on the total number of tokens in the sequences rather than using a fixed number of sequences per batch. This method permits for larger batches when processing many short sequences and restricts the batch to a single sequence when handling very long ones, thereby optimizing GPU memory utilization. We are currently using the implementation by *Rains* [24].

#### 2.6.3. Long sequences

For long sequences that cannot be processed individually (even with a batch size of one) without encountering out-of-memory errors, we employ a segmentation and pooling strategy to prevent truncation. Specifically, sequences are divided into overlapping chunks using a sliding window approach, and mean pooling is applied to generate the final embedding representation.

Figure 6 demonstrates the efficacy of our approach for handling long sequences by comparing the cosine similarity between true embeddings and those generated using the segmentation and pooling method across 8393 genes from a single genome.

#### 2.6.4. Measures for clustering and probability calculations

All three clustering algorithms (DBSCAN, HDBSCAN, and our weighted multi-threshold single-linkage clustering) utilize a similarity matrix, the product of the approximate nearest neighbor search. This step significantly accelerates the process and minimizes memory consumption for the subsequent clustering step by reducing the number of pairwise comparisons required.

To optimize the efficiency of the weighted multi-threshold single-linkage clustering, we employ a Union-Find (UF) data structure for organizing, merging, and searching within clusters. Common functions are Just-In-Time (JIT) compiled using Numba, substantially reducing the execution time for frequent UF operations. Finally, probability calculations are accelerated via vectorized operations in NumPy. Combined, these measures significantly decrease the total runtime, enabling the analysis of large-scale datasets. The expected time for clustering and probability calculation, with 350434 embeddings of size 1024 (the OrthoDB dataset described in Section 2.8.1), with τ = 25, is approximately 75 seconds using the hardware specified in Table 5

### 2.7. Statistics and Output

The primary output is a CSV file where each row represents a protein, including the following annotations:

- Pangenomic classification for each protein (i.e., *core, shell*, or *cloud* ).
- Cluster ID classification for each protein.
- For weighted multi-threshold single-linkage clustering (Section 2.5.8), the probabilities for both the pangenomic category and cluster ID are reported, calculated as described in Equations 4 and 5.

Additionally, users can choose to include the following optional information:

- Heaps’ Law statistics (see Section 2.7.1).
- Phylogenetic-like analysis of final clusters (see Section 2.7.2).

#### 2.7.1. Heaps’ Law

Heaps’ Law describes the growth dynamics of the pangenome by modeling how the total number of unique protein clusters scales with the number of genomes analyzed. We apply two complementary power-law models to characterize pangenome openness: *Pangenome accumulation model* which is the total pangenome size P (N ) as a function of the number of genomes N follows:

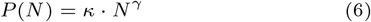

where κ is a constant and γ is the growth exponent. When γ > 0, the pangenome is considered *open*, indicating that new gene families continue to be discovered as more genomes are added. When γ < 0, the pangenome is *closed*, approaching a constant size asymptotically [25, 26]. The other power law is *New gene discovery model* which describes the rate at which new genes are discovered, Δ(N ), follows a complementary power law:

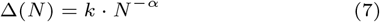

where k is a constant and α is the decay exponent. Here, α < 1 indicates an open pangenome (new genes continue to be discovered), while α > 1 indicates a closed pangenome (new gene discovery decays rapidly) [25, 26].

These two models are mathematically related through α = 1 − γ . Both formulations provide consistent assessments of pangenome openness, with the γ parameter directly indicating growth dynamics and the α parameter describing the decay in novel gene discovery rate. To estimate these parameters, we performed 300 random permutations of genome addition order and fitted the power-law functions to the averaged accumulation curves, ensuring robust statistical inference of pangenome characteristics.

#### 2.7.2. Phylogenetic-like Analysis

To generate a phylogenetic-like representation of the pangenome, it is possible to do the same clustering algorithm described in Algorithm 1 applied to the cluster centroids. Essentially, it builds a new cluster which effectively maps the relationships among clusters across the pangenome. The rationale for this phylogenetic-like analysis is to retrieve information about the relative (possibly evolutionary) positions of gene groups within the embedding space, potentially revealing biologically relevant features, such as identifying protein groups that are notably isolated. This approach introduces a clear bias which is that the diversity of protein clusters is represented as the cluster centroid, which is an oversimplification. The output of this phylogenetic-like analysis mimics that of the full method, as specified in the above subsection 2.7.

### 2.8. Datasets for evaluation

To address the challenge of evaluating pangenomic clustering software without a universal ground truth, we have chosen two datasets for the evaluation: a semi-synthetic dataset based on the OrthoDB v12 database and a experimental validated high-quality dataset from the CAFA 5 competition. Both datasets focus on the same task of validating protein clusters as a proxy for pangenomic classification.

#### 2.8.1. OrthoDB dataset creation

To evaluate our method on a controlled and biologically coherent dataset, a semi-synthetic pangenome was constructed from the OrthoDB v12 database [27, 28]. All organisms belonging to Bacteria (NCBI taxid: 2) were selected using the ‘odb12v1_level2species.tab’ file, and from this pool 100 organisms were randomly sampled with their corresponding protein sequences extracted from the master file ‘odb12v1_og_aa_fasta’ file. Following extraction, annotations were added via a multi-step pipeline: for each protein header, the OrthoDB orthologous group (OG) ID (from ‘odb12v1_OG2genes.tab’), functional cross-references such as GO terms and InterPro domains (from ‘odb12v1_gene_xrefs.tab’), and a protein product description (from ‘odb12v1_genes.tab’) were included; proteins labeled “hypothetical protein” or lacking a product description were removed, yielding a final dataset of 350434 proteins across 100 bacterial organisms.

##### OrthoDB dataset drawbacks

It should be mentioned that OrthoDB uses sequence alignment using MMseq2 as its method to define homologs/orthlogs [29, 27]. As mentioned in the introduction this potentially have some drawbacks, in relation to detection of homology and gene sematic. Furthermore, this dataset may introduce a bias toward algorithmic concordance (that is, model-inferred labels), possibly at the expense of biological concordance. Nevertheless, OrthoDB remains a relevant benchmark given the historical and ongoing use of sequence alignment for protein comparison.

#### 2.8.2. CAFA 5 dataset

To counterbalance the biases of the OrthoDB dataset, evaluation was expanded to include the CAFA 5 dataset. The CAFA 5 dataset consists of only experimentally determined GO terms for each protein. The dataset was created from UniProtKB in late 2022 and early 2023 and consists of 142246 sequences with all associated GO terms. The dataset consists of three subcategories of GO terms: Molecular Function (MF), Biological Process (BP), and Cellular Component (CC). This dataset must be viewed as a ground truth or gold standard for experimental validation, intended to counterbalance the biases found in the OrthoDB dataset.

### 2.9. Evaluation

As mentioned in 2.8, evaluating pangenome clustering is challenging due to the lack of a universal ground truth for pangenomic classes. To establish a robust benchmark, this study employs two distinct evaluation strategies: cluster quality and term consistency. Both strategies leverage the hierarchical information inherent in OrthoDB orthologous groups and GO terms.

#### 2.9.1. Cluster quality

To evaluate cluster quality we use Information Content (IC)[30, 31], a metric to measure the specificity of terms within a taxonomy. The IC of a given term is inversely proportional to its frequency, which means that rare and highly specific terms carry a higher IC value and vice versa.

IC calculation begins with determining the probability of a term’s t occurrence denoted as p(t). p(t) is calculated as the frequency of that term and all of its descendants divided by the total number of unique proteins in the corpus:

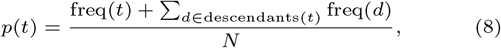

freq(d) is the number of genes annotated with a given term t, the summation accounts for all annotations to its descendants, and N is the total number of unique proteins annotated in the datasets 2.8.1 and 2.8.2. From this probability, the Information Content for term t is formally defined as its negative log-likelihood (Equation 9). This transformation ensures that less frequent, more specific terms yield a higher IC value.

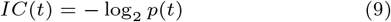

With IC defined, we can assess the quality of individual clusters. The Semantic Similarity Score measures cluster homogeneity by finding the Most Specific Common Ancestor (MSCA) of all genes within a cluster in a given hierarchy and reporting its IC value. Therefore, a higher score indicates a more specific, coherent cluster. For the semi-synthetic dataset, this score is calculated against both the OrthoDB identifier and GO hierarchies.

#### 2.9.2. Functional consistency

Solely for the semi-synthetic OrthoDB dataset, it is also possible to measure the homogeneity of gene product annotations. The score is the percentage of genes whose product description matches the cluster’s most frequent (dominant) annotation, after excluding non-informative terms.

#### 2.9.3. Term consistency

In some contrast but related to cluster quality, described in Section 2.9.1, we also evaluate the consistency of terms being clustered together using Term Consistency Score (TCS). The process begins by calculating a consistency score for each individual GO term, t. This score rewards functional specificity IC(t) while penalizing the term’s fragmentation across multiple clusters (Nc(t)), as shown in Equation 10.

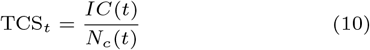

The final, overall TCS is then calculated as a weighted average of these individual term scores. The weight for each term is its relative frequency among all annotations in the dataset, which we denote as taf(t) (term annotation frequency). Meaning that more frequently annotated terms result in a higher weight for that term.

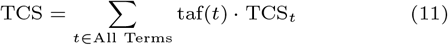

## 3. Results

In this work, we introduce a novel approach to pangenomic analysis using pLM encoders, which transform protein sequences into high-dimensional vectors, embeddings.

Additionally, we propose a new weighting scheme for multi-threshold single-linkage clustering method. To facilitate this, we have developed a comprehensive, user-friendly pipeline (implemented as a Python package) that streamlines the entire workflow, making complex computational tasks accessible. This framework aims to supports various pLM architectures for computing embeddings (for now only ESM2 3B and ProtT5-Xl), enabling users to conduct pangenomic analyses using diverse pretrained models and objectives.

### 3.1. Results for OrthoDB and CAFA5

On the OrthoDB label inferred via sequence alignment dataset (in Section 2.8.1), our proposed method resulted in clusters that have higher IC scores, characterized by higher functional specificity and homogeneity compared to SCARAP. There is a marked increase in values for both Mean Hierarchical Purity and Mean GO Coherence; for instance, the ProtT5 DBSCAN configuration achieved a Mean GO Coherence of 6.36, almost double that of SCARAP (3.59). Similarly, Mean Functional Consistency (FC) reached 99.82% with ESM2 3B Multi Threshold, significantly surpassing SCARAP’s 84.01%. Conversely, the Term Consistency Score revealed a trade-off in performance. SCARAP achieved a superior TCS (0.87) compared to the best performing configuration of the proposed method (0.63 for ESM2 3B Multi Threshold). This dissociation implies that the proposed method prioritizes the generation of highly functionally pure and coherent clusters, whereas SCARAP emphasizes the consolidation of terms into broader groups, potentially at the cost of internal consistency.

As discussed in Section 2.8.1, the generation of ground truth labels for the OrthoDB dataset relies on MMseqs2. Since SCARAP operates also using MMseqs2 algorithmic concordance likely introduces a bias that favors SCARAP. Consequently, the observed performance advantage may reflect model-inferred similarities rather than purely biological discordance.

On the CAFA 5 experimental validation dataset (Section 2.8.2), the proposed method demonstrates superior results compared to SCARAP across nearly all metrics, as illustrated in Table 3. Specifically, the proposed method produces clusters that are significantly more functionally consistent, often surpassing SCARAP by a two-fold margin in Weighted Mean GO Coherence (e.g., 3.54 for ProtT5 HDBSCAN vs. 1.53 for SCARAP). Regarding Term Consistency Score, while SCARAP excelled in the sequence-based OrthoDB dataset, it maintains a slight advantage on the experimental CAFA 5 dataset. Here, SCARAP (0.1380) gets a similar score to using ESM2 3B embeddings with Multi-Threshold clustering (0.1374). SCARAP is performing slightly better in TCS than all combinations of our method.

**Table 3.** Performance comparison on the CAFA5 dataset. The best performance for each metric is highlighted in bold. Default settings were used for SCARAP. Parameters for the weighted multi-threshold single-linkage clustering are model-specific: for ProtT5, mean = 0.856 and SD = 0.1259; for ESM2 3B, mean = 0.9935 and SD = 0.0076. For density-based clustering ((H)DBSCAN), parameters were set to ϵ = 0.05 and minimum cluster size = 2 for ESM2 3B model. For ProtT5 the parameters for density-based clustering was set to ϵ = 0.045 and minimum cluster size = 2. FC denotes Functional Consistency.

| Metric | ProtT5 Embeddings |  |  | ESM2 3B Embeddings |  |  | SCARAP |
| --- | --- | --- | --- | --- | --- | --- | --- |
|  | Multi Threshold | DBSCAN | HDBSCAN | Multi Threshold | DBSCAN | HDBSCAN |  |
| Mean GO (BP) | 5.8339 | **5.9612** | 5.6413 | 5.6193 | 5.6430 | 5.5095 | 4.3721 |
| Weighted Mean GO (BP) | 2.3396 | 2.5247 | **3.5395** | 2.1386 | 2.1386 | 2.5196 | 1.5257 |
| Mean GO (MF) | 4.1111 | 4.2334 | 4.0913 | 4.2759 | **4.2835** | 4.1845 | 2.8506 |
| Weighted Mean GO (MF) | 1.9576 | 2.1079 | **2.9253** | 1.9305 | 1.9305 | 2.2378 | 1.3211 |
| Mean GO (CC) | 3.3991 | **3.4551** | 3.29 | 3.2738 | 3.2819 | 3.1856 | 2.7359 |
| Weighted Mean GO (CC) | 1.4594 | 1.5644 | **2.1729** | 1.3062 | 1.3062 | 1.5265 | 1.0622 |
| Term Consistency Score | 0.1081 | 0.1016 | 0.1072 | 0.1374 | 0.1279 | 0.1309 | **0.1380** |

This shift in performance highlights a critical generalization gap for all evaluated methods. Transitioning from the semi-synthetic OrthoDB dataset to the experimentally validated CAFA 5 dataset results in a universal decline in TCS. SCARAP experiences a performance reduction of approximately 84.2% (from 0.8731 to 0.1380). The proposed method also experiences a significant, albeit slightly smaller, proportional reduction; for example, the ESM2 3B HDBSCAN configuration shows a drop of 76.5% (from 0.5573 to 0.1309). This universal decline suggests that transitioning from model-inferred labels to biologically verified functions introduces substantial complexity that challenges both sequence-identity and embedding-based clustering approaches. Crucially, because SCARAP and the proposed method converge to achieve effectively similar TCS scores on the CAFA 5 dataset, the proposed method’s substantial outperformance across all remaining metrics of functional consistency and coherence becomes the defining advantage on experimentally validated data.

### 3.2. Computational efficiency

Computational efficiency is important for large-scale pangenomics. In this regard, SCARAP is the sole winner, outperforming our method by a many-fold margin (around 11 folds). A runtime comparison is provided in Table 4, and the hardware specifications used for these computations are detailed in Table 5 in appendix. Table 4 highlights the computational efficiency of the models by comparing their embedding generation times, revealing that ESM2 3B is faster than the ProtT5 encoder. Furthermore, it details the respective execution times for the three clustering algorithms.

**Table 4.** Runtime (in seconds) across various embeddings, clustering algorithms, and the SCARAP pipeline. The CAFA5 dataset was used to generate the reported runtimes. The specifications of the machine used are described in the table 5 in appendix.

| Metric | ESM2 3B Embeddings | ProtT5 Embeddings | Multi-threshold | DBSCAN | HDBSCAN | SCARAP |
| --- | --- | --- | --- | --- | --- | --- |
| Time (s) | 4080 | 3520 | 50.8 | 27.1 | 79.5 | 324 |

**Table 5.** Server System Specifications.

| Component | Model / Type | Key Specifications |
| --- | --- | --- |
| CPU | Intel(R) Xeon(R) Silver 4410Y | Cores: 12<br>Threads: 24 |
| Memory (RAM) | DDR5 ECC Registered | Capacity: 128 GB |
| GPU | Nvidia A100 PCIe 80GB | GPU Memory: 80 GB HBM2e |

### 3.3. Pangenomic analysis on Streptomyces

Recently, 1034 high-quality actinomycete genomes were published [32]. These will act as the input for a large pangenomic analysis. While classical pangenomics typically focuses on a single species, this task expands the analysis to a higher taxonomic level, utilizing 1034 genomes from across the class *Actinomycetia*. The largest classical pangenomic analysis of *actinomycetes* that has been published uses 205 genomes [33].

Figure 4 shows that the pan-genome of the 1034 actinomycete genomes is open, with a γ value of 0.673. An open pan-genome indicates that the number of protein clusters continues to grow as more genomes are discovered and sequenced. This can be clearly observed in part A of Figure 4, where the relationship between the number of genomes and the number of protein clusters is illustrated, with no signs of the curve converging. This aligns well with the findings of the study on 205 genomes, which also demonstrated an open structure [33]. Part B of Figure 4 illustrates the relationship between the number of genomes and the proportions of pangenomic classes. The open pangenome is also evident from the convergence of all pangenomic classes, with the proportion of almost all proteins assigned the cloud label. The threshold values used are those explained in Table 1, the encoder model used is prot T5 and clustering method used is weighted multi-threshold single linkage clustering.

**Fig. 4.**
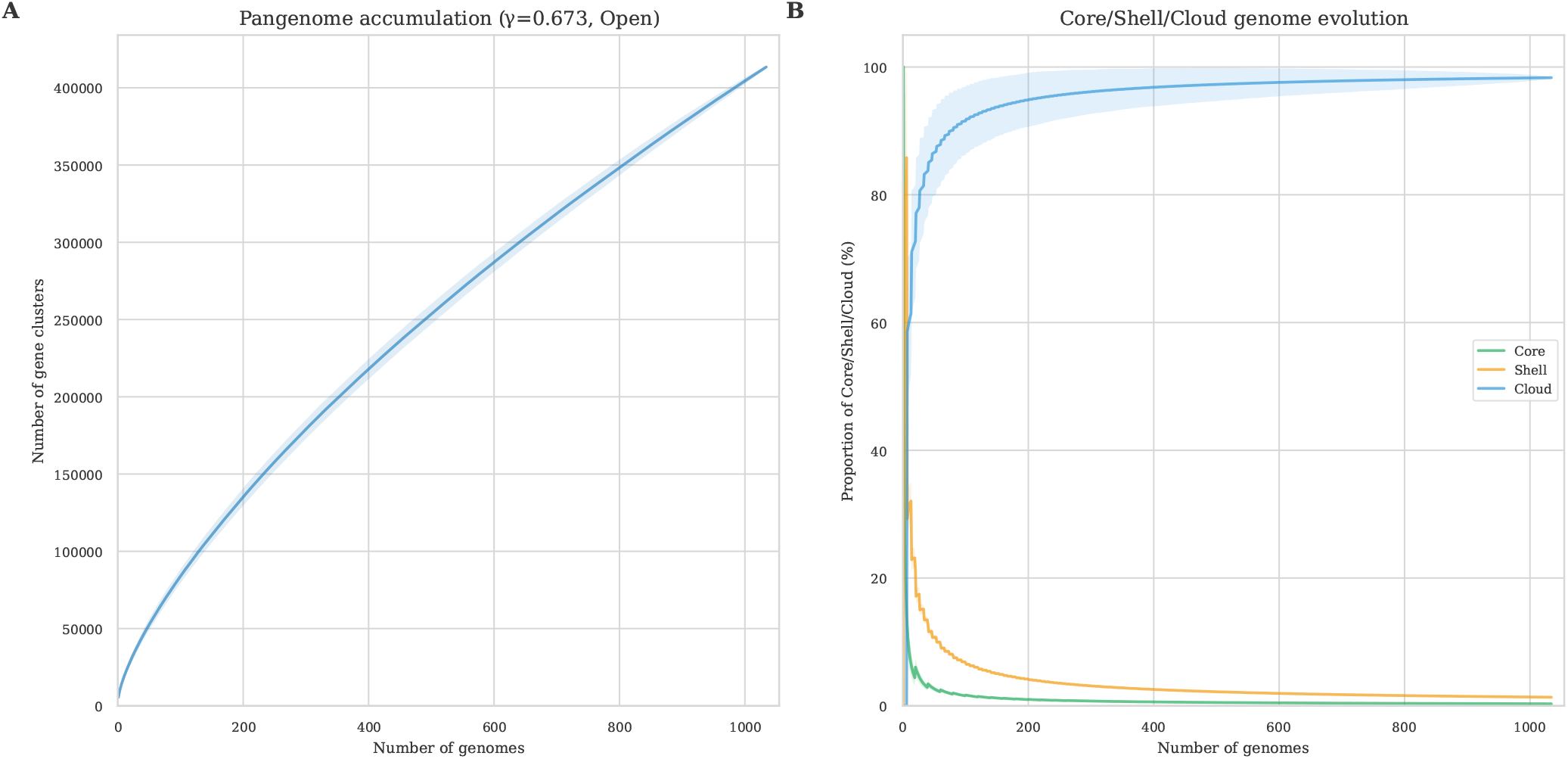
The data used for both parts of the plot is 1034 actinomycete genomes [32]. Part **A** of the figure illustrates the relation between number of genomes and number of protein clusters. It can be observed just by looking at the graph, that the group of actinomycete genomes displays an open genome. The reported gamma value of 0.67 is based on the equation 6. Part **B** of the plot illustrates the relation between number of genomes and the proportion of pangenomic classes, based on the thresholds in 1. What can be observed is the same effect as in part A of the plot, which shows that the group of actinomycete genomes displays an open pangenomic structure; this can be seen by the fact that the cloud proportion converges close to 100% percent, which means that the expectation is to find cloud genes when more genomes are added

It should be noted that part **B** of Figure 4 is the pangenomic proportions calculated using the number of protein clusters assigned to the pangenomic classes. This stands in contrast to using the number of individual proteins assigned to the respective pangenomic classes. The latter would result in larger core and shell proportions, due to the fact that these protein clusters have, by definition, more proteins in them.

Figure 4 can be saved if the corresponding flag is specified when the tool is run.

## 4. Discussion and Conclusions

Traditional pangenome construction relies on sequence alignment, which struggles to classify remote orthologs within the sequence identity “twilight zone”. To overcome this, our method leverages protein language model embeddings to capture deep structural and functional relationships beyond primary sequence similarity. Because a single global similarity threshold for single linkage clustering fails to capture the evolutionary diversity of different orthologous groups existing, we implemented a weighted multi-threshold single-linkage clustering approach. This applies a probabilistic weighting scheme derived from the normally distributed tipping points where orthologs transition from core to shell categories. While sequence-alignment tools like SCARAP perform well on the semi-synthetic OrthoDB dataset, this is likely due to an algorithmic concordance with the MMseqs2 tool used to generate the dataset’s labels. This advantage is largely nullified on the experimentally validated CAFA 5 dataset, where SCARAP experiences an 84.2% reduction in its Term Consistency Score, converging to a performance level effectively identical to our pLM-based approach. Crucially, because the TCS scores converge, the defining advantage of our method on experimental data lies in its superior cluster quality. Our method significantly outperforms SCARAP across all remaining metrics of functional consistency and coherence on the CAFA 5 dataset, achieving up to a two-fold increase in Weighted Mean GO Coherence. This demonstrates that the pLM-based approach offers substantially greater robustness and generalization when grouping biologically verified functions. Finally, we validated the scalability of our full method by characterizing 1,034 Actinomycetia genomes. The method successfully identified an open pangenome structure with a growth exponent of 0.67. Analysis of these genomes demonstrates that the method is practical for large datasets.

## 5. Acknowledgements

N.J.L, K.B. and P.C. have been supported by the Novo Nordisk Foundation (NNF25SA0109652).

## 6. Appendix

### 6.1. Weighted multi-threshold single linkage clustering

To address the heterogeneity of ortholog divergence where a single global threshold compromises between over merging and over splitting (Figure 3). Algorithm 1 executes a simultaneous sweep across a set of similarity thresholds {τ_1_, τ_2_, … }. Leveraging ANN search for efficiency, the method populates distinct Union-Find structures for every τ where pairwise similarity s ≥ τ . This procedure captures the granular transition from large clusters to singletons (Appendix Figure 5), generating the ensemble partitions required for the probabilistic categorization described in Section 2.5.8.

**Fig. 5.**
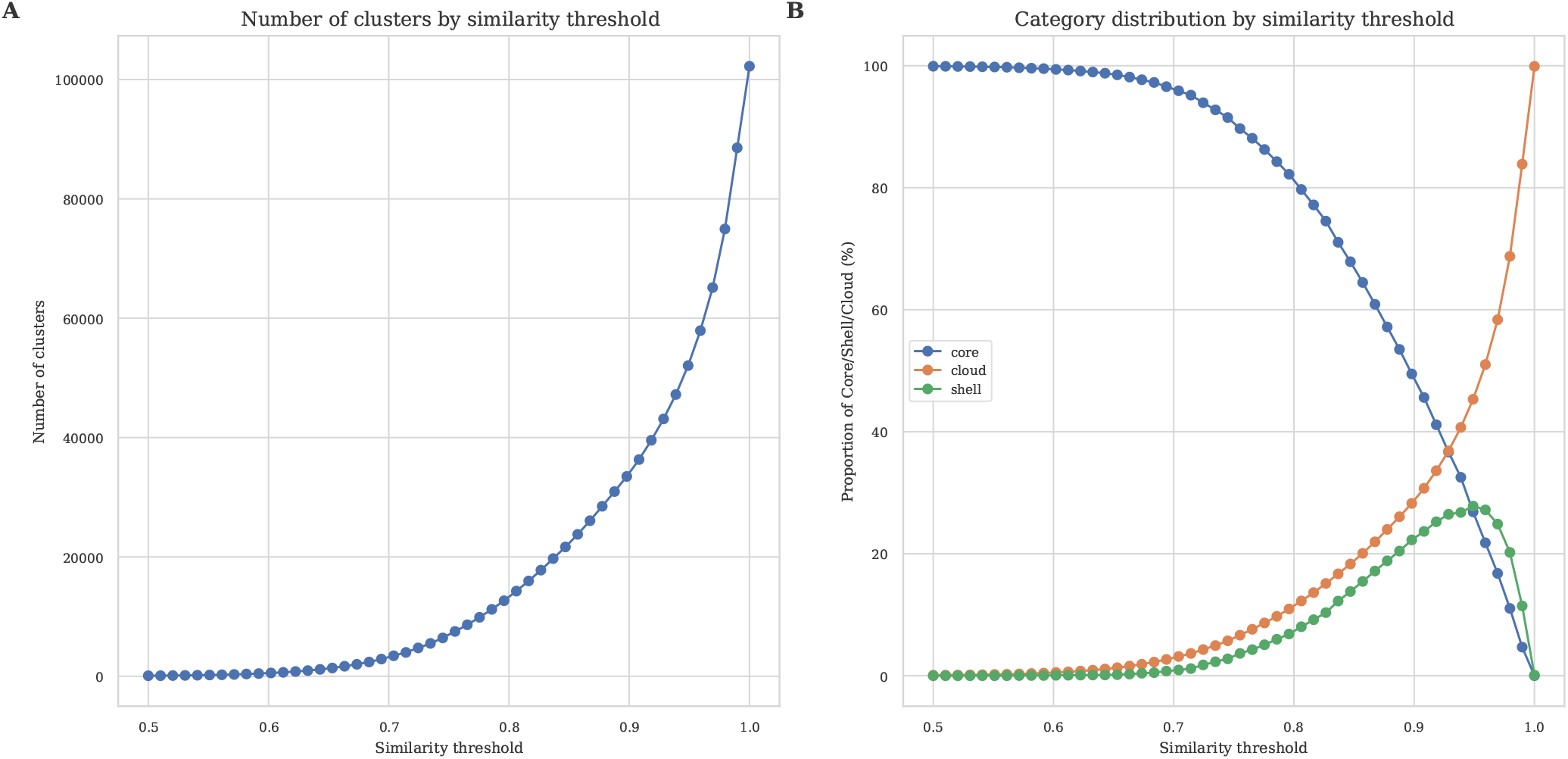
The **A** part of the figure, illustrates the relationship between the similarity threshold and number of clusters, intuitively the higher the similarity threshold the higher the number of clusters. The part **B** illustrates the relationship that similarity threshold has on the formation of pangenomic classes. The pattern shown seems to be very common, maybe even universal. The data used is a subset of streptomyces genomes [32]

#### Algorithm 1 Weighted multi-threshold single linkage clustering

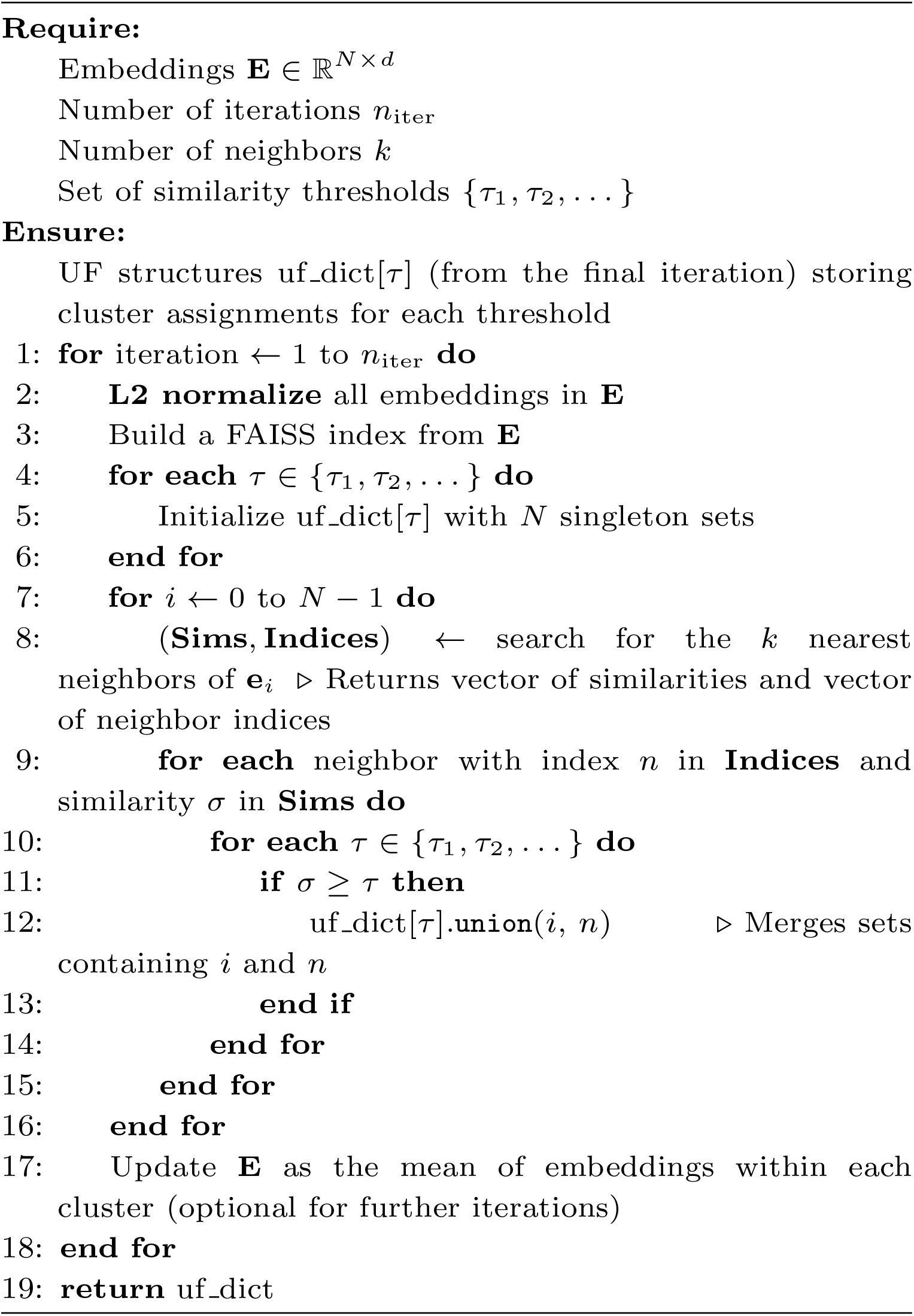

### 6.2. Supplementary Figures and Tables

**Fig. 6.**
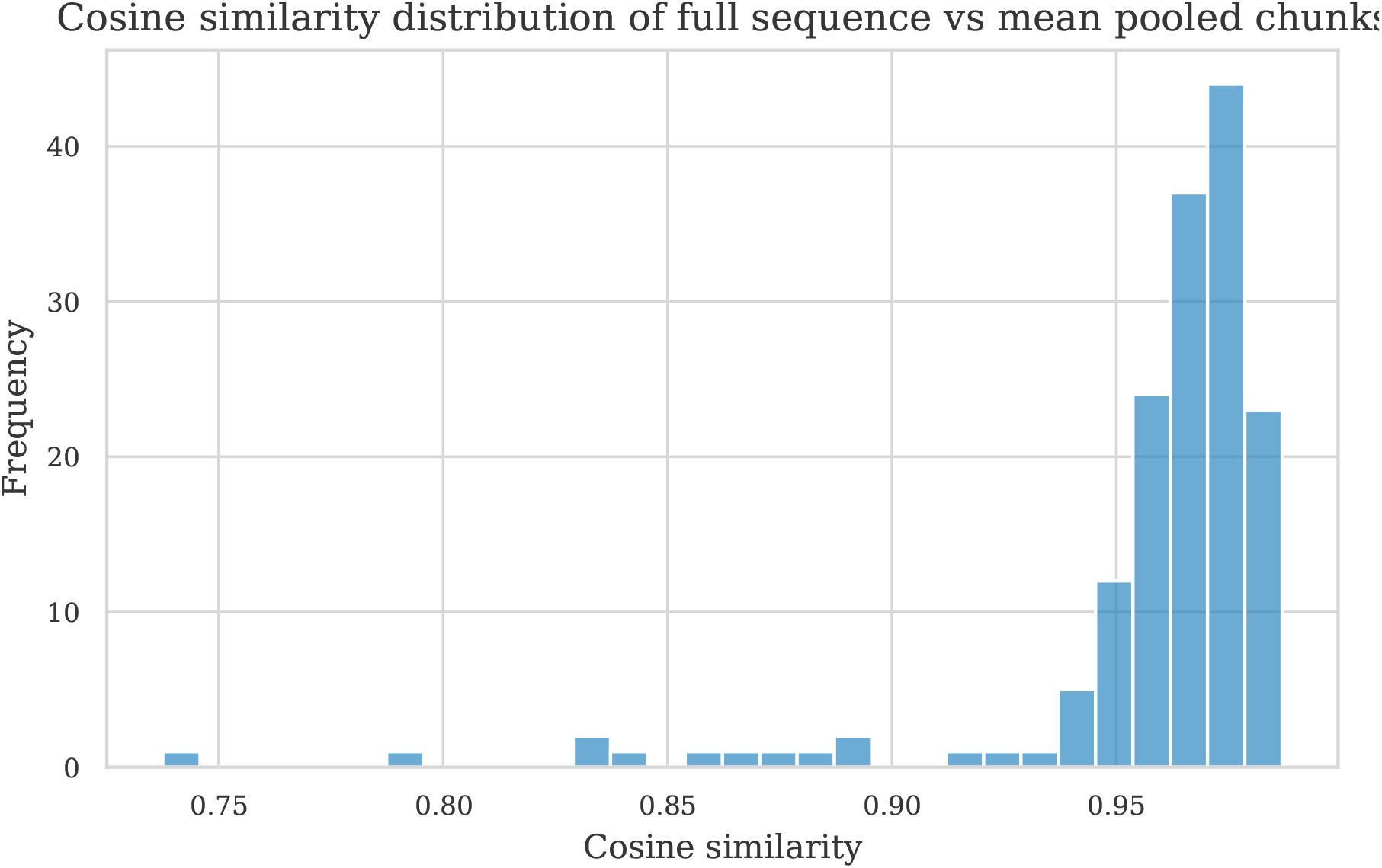
This figure illustrates that for very long input sequences, which cannot be processed on a GPU due to attention memory complexity, it is possible to chunk up the sequences and embed the chunks, then perform mean pooling of the chunks. This mean-pooled embedding is fairly similar to what can be observed if the full sequence is embedded.

**Fig. 7.**
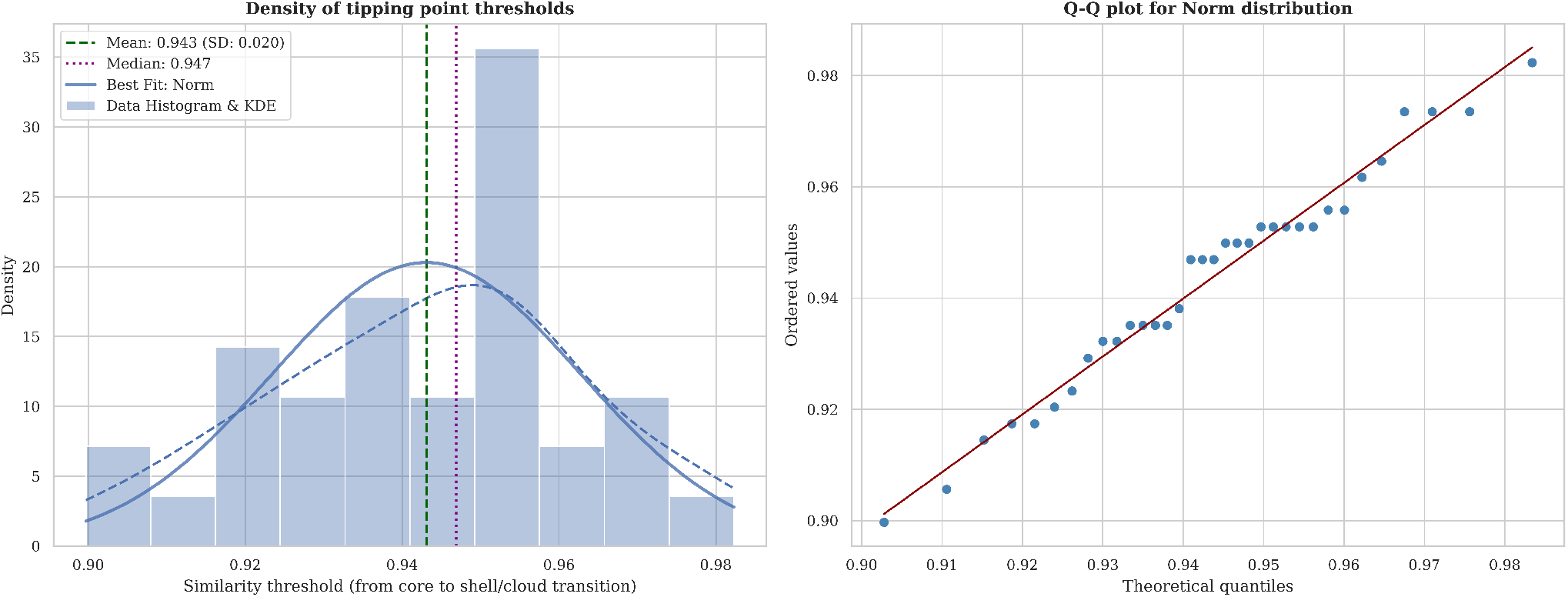
The figure illustrates that the density of the tipping points (core to shell transitions) of orthologous groups from OrthoDB forms a normal distribution. The transition from core to shell the core pangenome is 95% conservation of genes in a single cluster in accordance to Table 1

